# Metabolic differences and differentially expressed genes between C57BL/6J and C57BL/6N mice substrains

**DOI:** 10.1101/2022.07.06.499024

**Authors:** Shino Nemoto, Tetsuya Kubota

## Abstract

C57BL/6J (B6J) and C57BL/6N (B6N) mice are the most frequently used substrains in C57BL/6 (B6) inbred mice, serving as physiological models for *in vivo* studies and as background strains to build transgenic mice. However, little attention has been paid to the phenotypic differences between B6J and B6N mice, and they have often been attributed to mutations in the nicotinamide nucleotide transhydrogenase (*Nnt*) gene, which was found only in B6J. Nevertheless, phenotypic differences between the two that cannot be explained by *Nnt* mutations alone, especially in metabolic traits, indicate the presence of genetic variants associated with metabolism other than *Nnt*. We aimed to identify these genetic differences between B6J and B6N mice. Our results provide insights into differentially expressed genes (DEGs) in adipose tissues, skeletal muscle, liver, hypothalamus, and hippocampus, and phenotypic differences in metabolic traits between B6J and B6N. B6J mice had significantly lower body weight than B6N mice, regardless of a normal or high-fat diet. Blood insulin levels in B6J mice were significantly lower than those in B6N mice and glucose levels during dietary obesity were higher in B6J mice. Metabolic assessments revealed greater physical activity, less food intake, and higher energy expenditure in B6J mice than in B6N mice. Among the DEGs that were highly expressed in B6J mice compared to B6N mice, three genes—insulin degrading enzyme, adenylosuccinate synthase 2, and ectonucleotide triphosphate diphosphohydrolase 4—and the DEGs that had lower expression in B6J mice compared to B6N mice—*Nnt*, WD repeat and FYVE domain containing 1, and dynein light chain Tctex-type 1—were overlapped in all seven tissues. Our study provides insights into DEGs between B6J and B6N, which will be useful for substrain selection for mouse experiments, avoiding erroneous experimental results, and reviewing the results of studies that have used B6J, B6N, or mixed substrains.

## Introduction

C57BL-derived inbred mouse B6 is currently the most frequently used laboratory animal and is a vital tool in various biomedical studies, with more than 20 different substrains. Of the 20 different substrains, B6J and B6N mice are the most commonly used, owing to their strain stability and ease of breeding. B6J was the first mouse substrain that had its genome fully sequenced [1]; thus, many transgenic mice, including those produced using the Cre-lox and FLP-FRT recombination systems, have been generated with a B6J background, and *in vivo* studies using B6J as physiological or pathological models have increased rapidly. However, the B6N substrain has recently become more common and standardized because it was used as the embryonic stem cell line for large-scale knockout generation and phenotyping projects (*e*.*g*., International Knockout Mouse Consortium, International Mouse Phenotyping Consortium, and NIH Knockout Mouse Project), generating more than 5,000 targeted mutant mouse lines that are currently available to researchers worldwide [2]. Thus, both the B6J and B6N mice substrains have become indispensable and exclusive materials. Nevertheless, researchers are indifferent to the genetic and phenotypic differences between the two substrains [3, 4] and generate mice with mixed substrains [5] or do not explicitly address the substrain distinction in their publications [6]. Moreover, these unaddressed differences can lead to confounding experimental outcomes or misinterpretations; therefore, comprehensive information about the properties of these mouse substrains needs to be ascertained.

To date, various genomic differences such as indels, structural variations, and single nucleotide polymorphisms (SNPs), as well as phenotypic differences between B6J and B6N, have been reviewed [3, 7], some of which are known to be associated with pathogenesis and require caution when conducting experiments in such areas. For example, ophthalmic problems have been known to occur in B6N mice [7, 8], regardless of genetic engineering [9, 10], as the substrain carries a single nucleotide deletion that causes a frameshift mutation and subsequent protein truncation in the crumbs cell polarity complex component 1 (*Crb1*) gene, which is associated with retinal degeneration. Therefore, in ocular research, it would not be appropriate to choose B6N as a background or *in vivo* model, crossbreed it with B6J, or compare it to B6J. Similarly, these substrain differences must be considered when selecting mouse substrains in metabolic studies.

Many phenotypic differences [11-24] between B6J and B6N, especially in metabolic traits such as glucose tolerance, insulin secretion, and body weight, have been attributed to loss of function mutations in the nicotinamide nucleotide transhydrogenase *(Nnt*) gene, including deletion of five exons and a missense mutation within the *Nnt* locus, which is harbored only in B6J mice [11,25,26]. However, conflicting data that do not correlate with *Nnt* mutations exist for differences in body weight between the two substrains, with reports of B6J mice weighing more than B6N mice [12-14, 27, 28], B6J mice weighing less than B6N mice [15, 29-32], and no difference in body weight between the two substrains [16, 17, 33]. In addition, conflicting data also exist regarding whether glucose tolerance and insulin secretion are impaired in B6J or B6N [12-19, 29, 31, 33]. These contradictory reports suggest that *Nnt* is not the only genetic variant that needs to be considered, and other unknown variants that affect metabolic traits in these substrains need to be ascertained.

In this study, we aimed to investigate the genetic causes of the phenotypic differences between B6J and B6N substrains. We report candidate genes that are presumed to harbor variants in either B6J or B6N, along with comparative data on the metabolic and behavioral characteristics of B6J and B6N. We show that B6J mice had lower body weight, adiposity, blood insulin levels, and food intake, and were more active than B6N mice. Our results give new insights into differentially expressed genes (DEGs) between B6J and B6N, which will be useful not only in the selection of substrains for mouse experiments but also in reviewing and reinterpreting the results of studies that have already used B6J, B6N, or mixed substrains.

## Materials and Methods

### Animals and experimental design

All experimental procedures were approved and performed in accordance with the Institutional Animal Care and Use Committee of the RIKEN Yokohama Campus. 7 week old male C57BL/6J and C57BL/6NCrl mice were purchased from the Oriental Yeast Company, Ltd. (Shiga, Japan) and were acclimated for 1 week and maintained on an alternating 12 h light/dark cycle at a temperature of 23 °C, with free access to food and water. After the acclimatization period, the mice were randomly divided into two experimental groups per strain (n=5 per group). Normal diet (ND) (CLEA Rodent Diet CE-2:12% calories from fat, 59.1% of calories from carbohydrates, 28.8% of calories from protein; CLEA Japan Inc., Shizuoka, Japan) or a high-fat diet (HF) (Clea High-Fat Diet 32:56. 7% of calories from fat, and 23.1% of calories from carbohydrates, 20% calories from protein; CLEA Japan Inc.) for 30 weeks. At the age of 20–30 weeks old, a metabolic assessment was performed. At the age of 38 weeks old, mice were euthanized under isoflurane anesthesia, blood was collected, and organs, including inguinal white adipose tissue (iWAT), epididymal white adipose tissue (eWAT), brown adipose tissue (BAT), skeletal muscle (muscle), liver, hypothalamus (Hyt), and hippocampus (Hic), were rapidly removed, weighed, submerged in RNAlater solution (Thermo Fisher Scientific, Waltham, MA, USA) at 4 °C for 20 h, and stored at –20 °C.

### Plasma parameters

Blood glucose levels were determined using a compact glucose analyzer (Glutest Sensor; Sanwa Kagaku, Nagoya, Japan). Plasma insulin (Morinaga Institute of Biological Science, Kanagawa, Japan), leptin (R&D Systems, Minneapolis, MN, USA), and adiponectin (Otsuka Pharmaceutical Co., Ltd., Tokyo, Japan) levels were measured using an ELISA kit. Plasma triglyceride (TG), total cholesterol (T-Cho), high-density lipoprotein cholesterol (HDL), non-esterified fatty acids (FFA), alanine aminotransferase (ALT), aspartate aminotransferase (AST), and lactate dehydrogenase (LDH) levels were measured using reagents from Wako Pure Chemical Industries, Ltd. (Osaka, Japan). All assays were performed according to the manufacturer’s instructions.

### Metabolic assessments

Oxygen consumption (VO_2_) and carbon dioxide exhalation (VCO_2_) were measured using an open-circuit metabolic gas analysis system connected directly to a mass spectrometer (ARCO-2000; Arco Systems Inc., Chiba, Japan). The mice were housed in individual acrylic chambers with free access to food and water. After 5 days of adaptation, VO_2_ and VCO_2_ were measured in individual mice for 1 min at 15 min intervals over a 7 day period with airflow at 0.3 L/min. Total energy expenditure, carbohydrate consumption, and fat consumption were computed based on VO_2_ and CO_2_ production. Locomotor activity was estimated based on the number of infrared beams broken in both the x- and y-directions using an activity monitoring system combined with a food intake recording system (ACTIMO-100M/MFD-100M; Shin Factory, Fukuoka, Japan).

### RNA sequencing

#### Tissue preparation and RNA isolation

Minced tissues were homogenized with Sepasol RNAI solution (Nacalai Tesque, Kyoto, Japan) using a TissueLyser LT instrument (Qiagen, Hilden, Germany) set at 50 strokes/s for 5 min. The homogenate from adipose tissues was centrifuged at 3000 × *g* for 10 min, and the bottom layer was transferred into a new tube to separate the fat from the upper layer. Chloroform was then added to the sample, and the vortexed sample was centrifuged at 14000 × *g* for 10 min to separate the RNA phase. The RNA phase was then transferred to a new tube and subjected to total RNA purification using QIAcube and the RNeasy kit (Qiagen). Quality analysis of RNA samples was performed using TapeStation (Agilent Technologies, Santa Clara, CA, USA) and RNA ScreenTape (Agilent Technologies).

#### Library construction and sequencing

Libraries were generated with the NEBNext Ultra RNA Library Prep Kit for Illumina (New England Biolabs, Ipswich, MA, USA). mRNA was enriched from total RNA (250 ng) using magnetic poly-T beads. First- and second-strand cDNAs were synthesized using random hexamer primers, M-MuLV reverse transcriptase, DNA polymerase I, and RNase H, followed by the conversion of overhangs to blunt ends. DNA fragments were ligated with NEBNext adaptors and size-fractionated with the AMPure XP system (Beckman Coulter, Inc., CA, USA) before treatment with the USER enzyme (New England Biolabs) and polymerase chain reaction (PCR) amplification with universal and index primers using Phusion high-fidelity DNA polymerase. PCR products were purified using the AMPure XP system, and the quality of the library was assessed using the TapeStation system (Agilent Technologies). Pooled libraries were sequenced on an Illumina HiSeq 2500 platform to obtain 50 bp single-end reads.

#### Read mapping and quantification of gene expression level

Reads were mapped to genes in the reference mouse genome (UCSC mm9) and assembled into transcripts, whose abundance was estimated as the expected number of fragments per kilobase per million base pairs sequenced (FPKM) using Cufflinks (v 1.3.0). Bowtie (v 0.12.7) was used to build an index of the reference genome and TopHat (v 1.4.0) was used to align the reads.

### Differential gene expression analysis

Data were analyzed using Strand NGS (v. 2.7, Strand Life Sciences, Bengaluru, India). DESeq1 was used to compare pairs of sample groups that included four biological replicates. P-values were adjusted using the method of Benjamini and Hochberg to control for the false discovery rate. The threshold for significant differential expression was set at q<0.05. The summarized data were then assessed using statistical models (Mann–Whitney unpaired test, Benjamini-Hochberg multiple gene correction) to generate gene lists of the DEGs.

### Statistical Analysis

Statistical analyses were performed using GraphPad Prism 8 software. Quantitative two-group data were analyzed using an unpaired two-tailed t-test. A comparison of data with two factors was performed using a two-way analysis of variance, followed by Tukey’s test. Statistical significance was set at *p*<0.05 or *p*<0.01.

## Results

### B6J mice have lower body weight than B6N mice

There were significant differences in body weight between the B6J and B6N groups, regardless of whether they were fed ND or HF (Fig 1). Regression analysis showed that B6J mice on ND and HF gained weight 2-fold [*y* = 6.4*Ln*(*x*) + 9.7 R^2^ = 0.99 vs *y* = 12.6*Ln*(*x*)− 3.2, R^2^ = 0.99] and 1.2-fold [*y* = 23.0*Ln*(*x*) − 24.0 R^2^ = 0.97 vs *y* = 26.7*Ln*(*x*) − 28.7, R^2^ = 0.97] less than B6N mice did, respectively.

**Fig 1.**
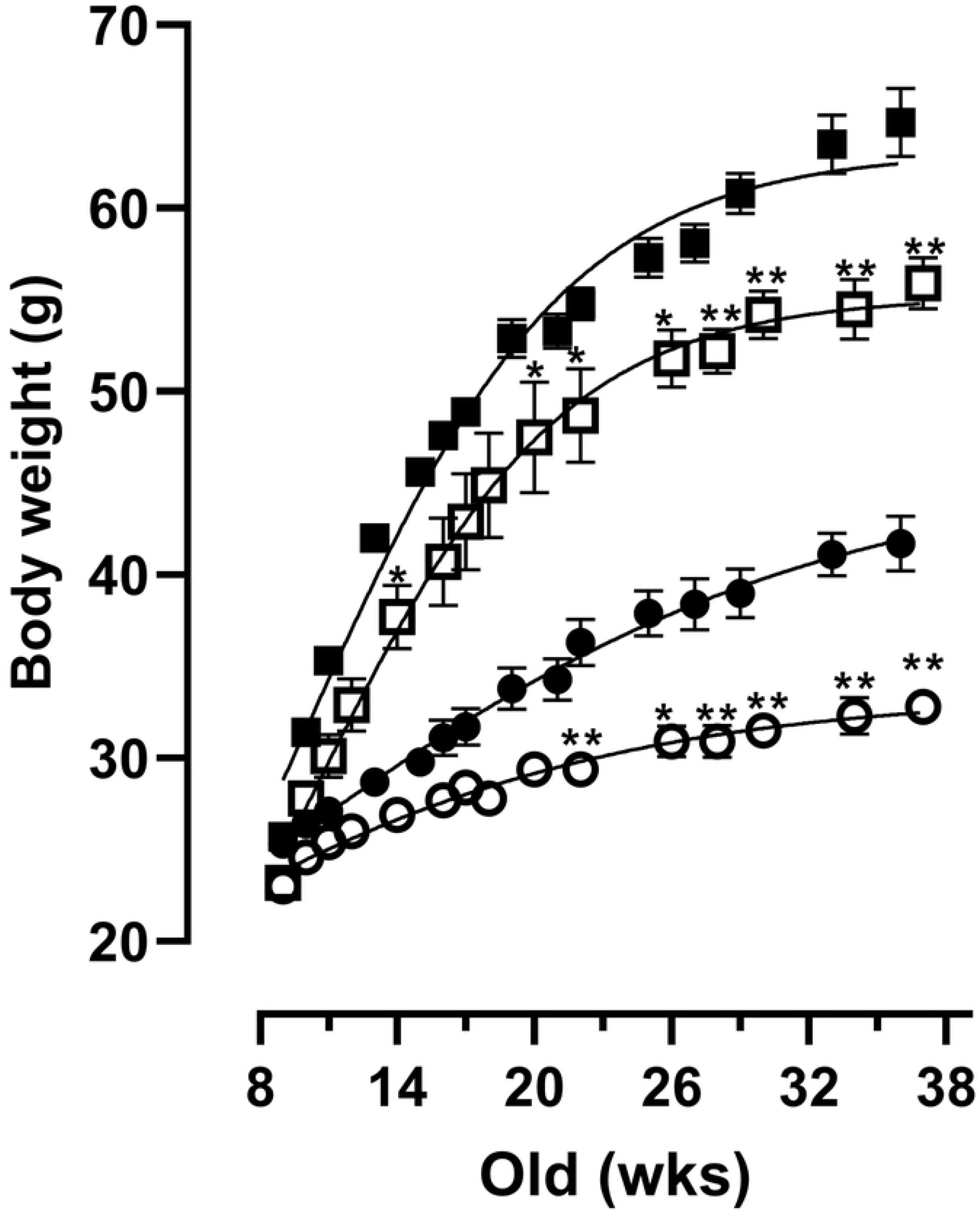
Difference in body weight between B6J and B6N mice on ND or HF. The body weight of male B6J and B6N mice on ND or HF was monitored up to 38 weeks old age (n = 5 per group). Data are presented as mean ± SEM. Asterisks (*) denote significant differences (\**p*<0.05, \*\**p*<0.01) between B6J and B6N mice at each time point in the same food group. ND, normal diet; HF, high-fat diet. Open circles, ND-fed B6J; filled circles, ND-fed B6N; open squares, HF-fed B6J; filled squares, HF-fed B6N.

### B6J mice exhibit lower adiposity than B6N mice

The weight of the white adipose tissues (iWAT and eWAT) of ND-fed B6J mice was significantly smaller than that of B6N mice, when measured in absolute weight (Fig 2A and 2B) and when normalized to body weight (Fig 2F and 2G). In the group of mice fed HF, the iWAT weight of B6J mice tended to be lower than that of B6N mice (Fig 2A and 2F), whereas eWAT was similar in terms of absolute weight (Fig 2B), and relative weight; however, eWAT weight was slightly higher in B6J mice (Fig 2G). Similarly, the BAT of ND-fed B6J weighed less whereas that of HF-fed B6J weighed more compared to that of B6N mice fed ND and HF, respectively (Fig 2C and 2H). Muscle size also tended to be smaller in the B6J group than in the B6N group in terms of absolute weight (Fig 2D). However, in terms of relative weight, it was significantly higher in B6J mice than in B6N mice in both the ND and HF groups (Fig 2I). Liver weight was slightly less in absolute weight in ND-fed B6J mice (Fig 2E); however, liver weight was slightly more in relative weight (Fig 2J) compared to B6N mice, and no significant difference was observed under HF conditions (Fig 2E and 2J). Looking at the effect of the high-fat diet on tissue weight, HF caused a significant increase in the liver in both B6J and B6N mice (Fig 2E) and in the iWAT and BAT weights of B6J mice but not B6N mice (Fig 2A and 2C). Surprisingly, the HF did not increase eWAT weight in both B6J and B6N mice; rather, eWAT weight in B6N mice was significantly less in the HF group than in the ND group (Fig 2B). Collectively, these results indicate that B6J mice have lower adiposity and more are susceptible to a HF than are B6N mice.

**Fig 2.**
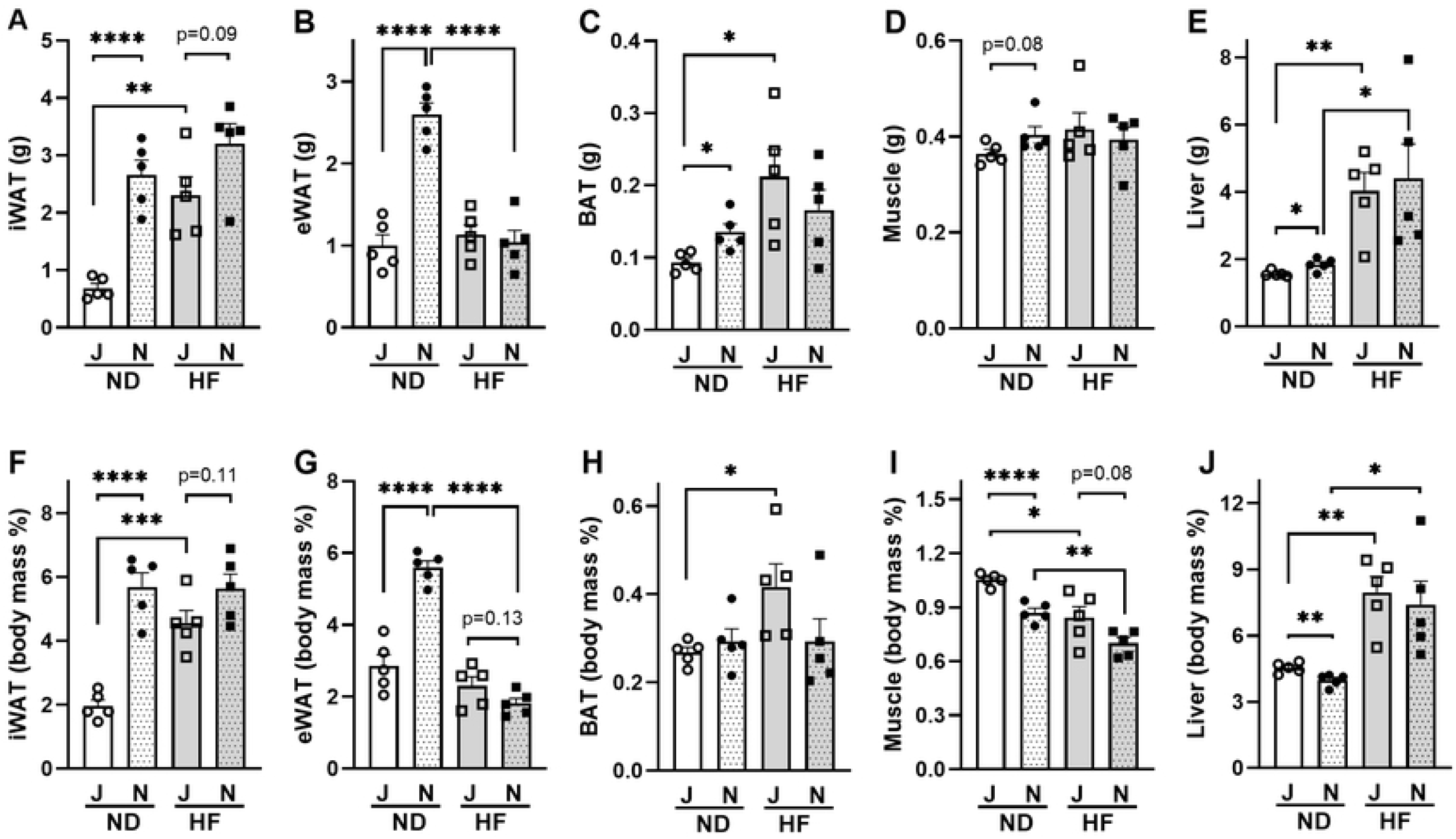
Tissue weights in B6J and B6N mice on ND or HF. (A–E) Absolute weight (gram) and (F–J) relative weight calculated by organ weight/body weight (body mass percentage). Values are means + SEM (n=5), and asterisks (*) indicate significant differences (\**p*<0.05, \*\**p*<0.01, \*\*\**p*<0.001, and \*\*\*\**p*<0.0001, two-way ANOVA, followed by Tukey’s test). ND, normal diet; HF, high-fat diet; iWAT, inguinal white adipose tissue; eWAT, epididymal white adipose tissue; BAT, brown adipose tissue; Muscle, skeletal muscle. Open circles, ND-fed B6J; filled circles, ND-fed B6N; open squares, HF-fed B6J; filled squares, HF-fed B6N.

### Plasma leptin and adiponectin levels in B6J mice are lower than those in B6N mice

To confirm the lower adiposity of B6J mice than B6N mice (Fig 2A and 2F), we measured two adipokines: leptin and adiponectin. In the ND-fed groups, the plasma leptin levels of B6J mice were significantly lower than those of B6N mice (Fig 3A). The adiponectin levels were also lower in ND-fed B6J mice than in ND-fed B6N mice but in a slightly weaker trend (*p* = 0.07, Fig 3B). In the HF-fed groups, adipokine levels were also lower in B6J mice than in B6N mice; however, the difference is not statistically significant (Fig 3A and 3B). Since leptin increases with fat accumulation and adiponectin decreases, the leptin/adiponectin ratio correlates well with adiposity [34]. In addition, this ratio is correlated with insulin resistance, which may represent adipose tissue dysfunction. Therefore, we calculated the ratio of leptin to adiponectin and found it to be approximately four-fold lower in B6J mice than in B6N fed ND (Fig 3C), confirming that the B6J substrain has lower adiposity than B6N.

**Fig 3.**
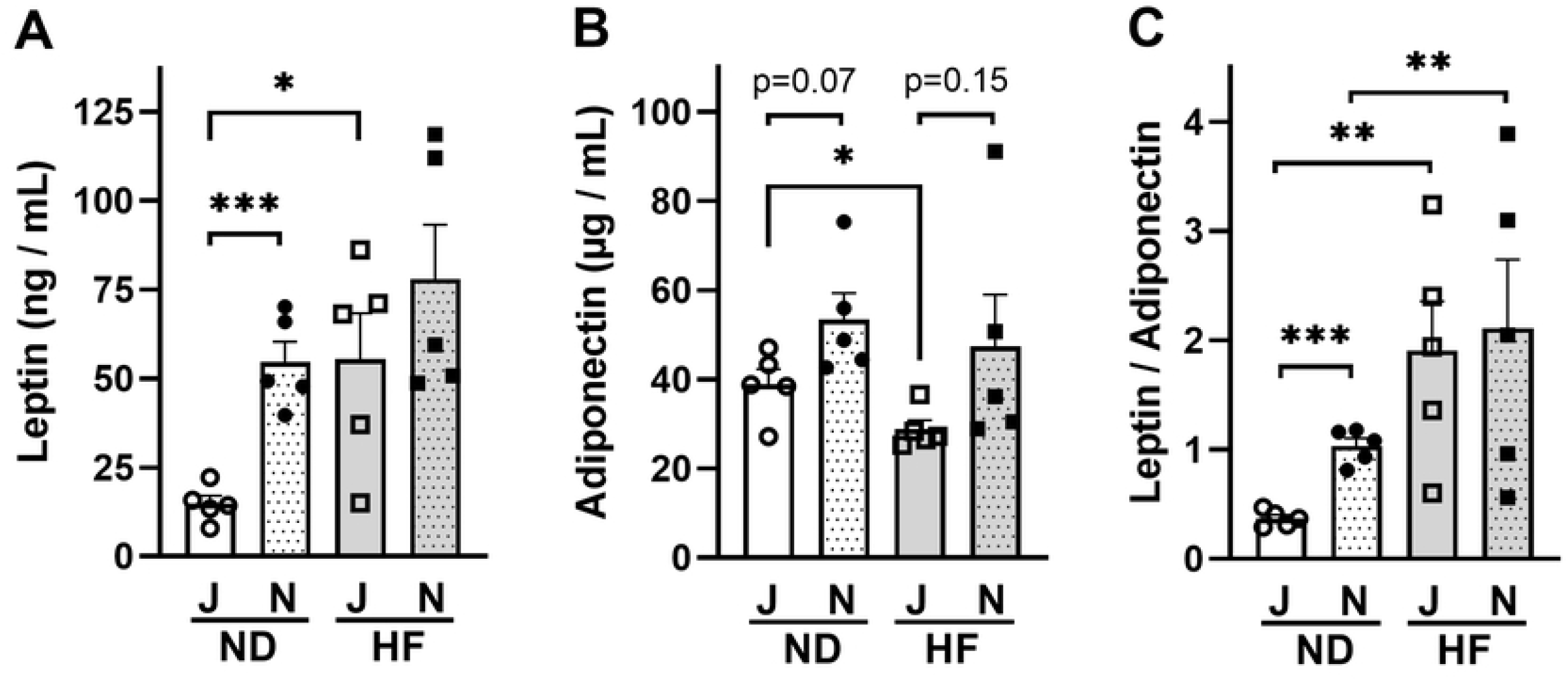
Differences in plasma leptin, adiponectin, and leptin to adiponectin ratio in B6J and B6N mice on ND or HF. (A) Leptin, (B) Adiponectin, (C) Leptin/Adiponectin. Values are means + SEM (n=5), and asterisks (*) indicate significant differences (\**p*<0.05, \*\**p*<0.01, and \*\*\**p*<0.001, two-way ANOVA, followed by Tukey’s test). ND, normal diet, HF, high-fat diet. Open circles, ND-fed B6J; filled circles, ND-fed B6N; open squares, HF-fed B6J; filled squares, HF-fed B6N.

### Plasma insulin levels of B6J mice are lower than those of B6N mice

We further compared commonly measured blood metabolic parameters, such as glycemic (glucose and insulin), lipidic (T-Chol, HDL, FFA, and TG), and liver enzymes (ALT, AST, and LDH). Insulin and FFA concentrations differed significantly between B6J and B6N mice (Fig 4A and 4B). Insulin concentrations were lower in B6J than in B6N under both ND and HF conditions (Fig 4A). FFA was significantly lower in HF-fed B6J mice than in HF-fed B6N mice (Fig 4B). Although T-Cho (Fig 4C), HDL (Fig 4D), ALT (Fig 4E), AST (Fig 4F), and LDH levels (Fig 4G) were similar between B6J and B6N mice, they were significantly higher in the HF groups than in the ND groups (Fig 4C–4G). Additionally, the blood glucose level was higher in the HF groups than in the ND groups for both B6J and B6N; however, between B6J and B6N mice, B6J mice showed a greater increase in glucose levels than B6N mice (Fig 4H). Similarly, TG concentrations during HF feeding were lower than those during ND feeding for both B6J and B6N mice; however, between B6J and B6N mice, B6J mice showed a greater decrease in TG concentration than B6N mice (Fig 4I).

**Fig 4.**
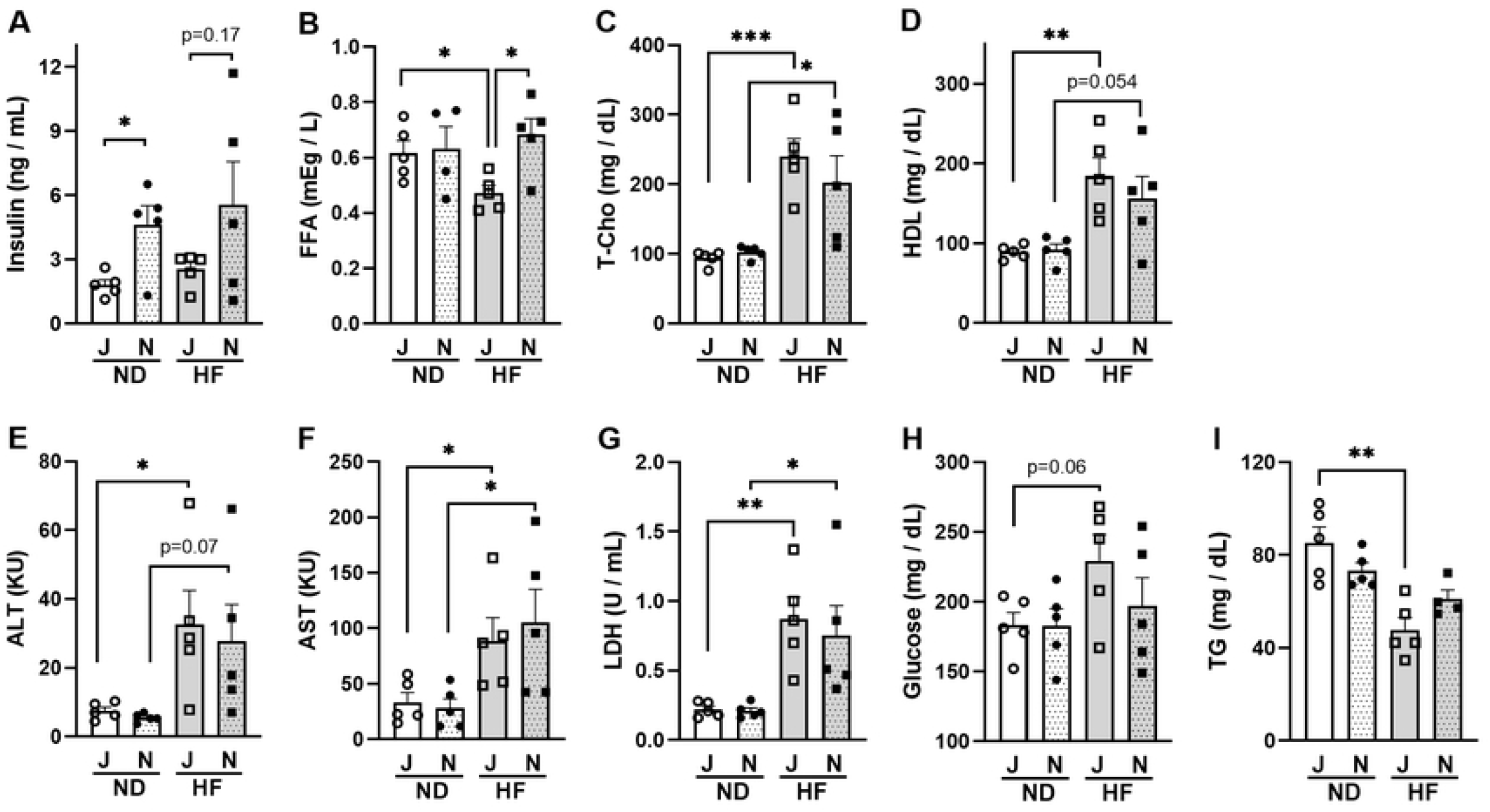
Differences in metabolic parameters in the plasma of B6J and B6N mice on ND or HF. (A) insulin, (B) FFA, non-esterified fatty acids, (C) T-Cho, total cholesterol, (D) HDL, high-density lipoprotein cholesterol, (E) ALT, alanine aminotransferase, (F) AST, aspartate aminotransferase, (G) LDH, lactate dehydrogenase, (H) glucose, (I) TG, triglyceride. Values are means + SEM (n=5), and asterisks (*) indicate significant differences (\**p*<0.05, \*\**p*<0.01, and \*\*\**p*<0.001, two-way ANOVA, followed by Tukey’s test). ND, normal diet, HF, high-fat diet. Open circles, ND-fed B6J; filled circles, ND-fed B6N; open squares, HF-fed B6J; filled squares, HF-fed B6N

### B6J mice consume more energy than B6N mice

To determine the difference in metabolic rates between B6J and B6N mice, we measured oxygen consumption and carbon dioxide production, and calculated energy, carbohydrate, and fat consumptions (normalized to body weight). Values are presented per diet and per light/dark phase (“Dark” for the active phase and “Light” for the inactive phase, “24 h” throughout both phases). Although the overall differences between B6J and B6N mice were not large, some significant differences were observed such as energy expenditure, with B6J mice tending to expend more energy than B6N mice during the active phase (Fig 5A). Similarly, there was a significant difference in carbohydrate consumption when fed ND, with B6J mice consuming more carbohydrates than B6N mice during the active phase (Fig 5B). Fat consumption data also showed differences between B6J and B6N when fed HF, with B6J mice consuming more fat than B6N mice throughout the day (Fig 5C). In addition, behavioral factors that affect metabolic rates, such as physical activity and food intake, were assessed. B6J mice tended to be more active than B6N mice when fed both ND or HF (Fig 5D) and took significantly less food (Fig 5E).

**Fig 5.**
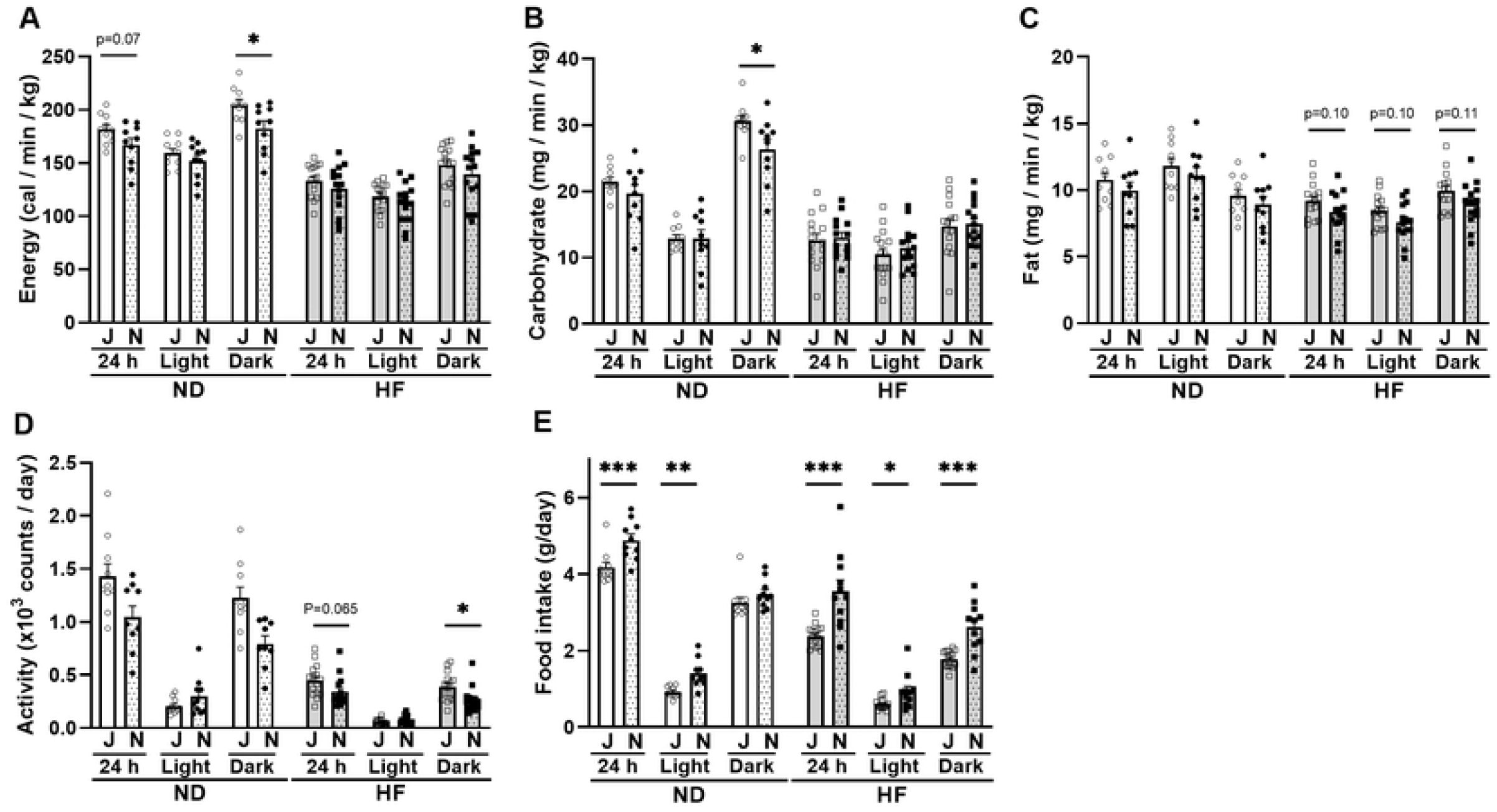
Differences in metabolic assessments between B6J and B6N mice. (A) Energy expenditure (B) Carbohydrate consumption (C) Fat consumption (D) Activity (E) Food intake. Values are means + SEM (n=5, experimented in duplicates), and asterisks (*) indicate significant differences (\**p*<0.05, \*\**p*<0.01, and \*\*\**p*<0.001). Open circles, ND-fed B6J; filled circles, ND-fed B6N; open squares, HF-fed B6J; filled squares, HF-fed B6N.

### DEGs between B6J and B6N

Table 1 shows the number of DEGs (*p* < 0.05) in the iWAT, eWAT, BAT, muscle, liver, hypothalamus, and hippocampus between B6J and B6N mice. Comparable numbers of DEGs were upregulated and downregulated in each tissue in the ND and HF groups, respectively. The gene names of the DEGs in each tissue are listed in S1 Table, along with their average fold change (FC), *p*-values, and false discovery rate (FDR). Among the DEGs that were highly expressed in B6J mice compared to B6N mice, three genes, insulin degrading enzyme (*Ide*), adenylosuccinate synthase 2 (*Adss2*), and ectonucleotide triphosphate diphosphohydrolase 4 (*Entpd4*), in the ND group and five genes, *Ide, Adss2, Entpd4*, B-TFIID TATA-box binding protein associated factor 1 (*Btaf1*), and transmembrane protein 267 (*TMEM267*), in the HF group overlapped in all seven tissues examined in the present study. Likewise, among the DEGs that showed lower expression in B6J mice compared to B6N mice, four genes, *Nnt*, WD repeat and FYVE domain containing 1 (*Wdfy1*), dynein light chain Tctex-type 1 (*Dynlt1*), and RAB4A, member RAS oncogene family (*Rab4A*), in the ND group and three genes, *Nnt, Wdfy1*, and *Dynlt1* in the HF group were identified. Of these overlapping genes, *Wdfy1* showed high FC and was significantly (FDR < 0.1) lower in ND-fed B6J mice by approximately 4- to 6-fold compared to B6N mice (Fig 6E). *Entpd4* also showed consistent FC in the HF group with approximately 2- to 3-fold higher expression in B6J mice than in B6N (Fig 6C).

**Table 1.** Number of differentially expressed genes (DEGs) in the iWAT, eWAT, BAT, muscle, liver, Hyt, and Hic between B6J and B6N mice fed a normal diet (ND) or high-fat diet (HF).

|  |  | J > N | J < N |  |  | J > N | J < N |
| --- | --- | --- | --- | --- | --- | --- | --- |
| iWAT | ND | 731 | 790 | Liver | ND | 972 | 629 |
|  | HF | 888 | 1349 |  | HF | 768 | 906 |
| | ND ∩ HF | 29 | 31 | | ND ∩ HF | 85 | 44 |
| eWAT | ND | 2450 | 753 | Hyt | ND | 1037 | 1987 |
|  | HF | 1190 | 2547 |  | HF | 2109 | 1163 |
| | ND ∩ HF | 160 | 105 | | ND ∩ HF | 186 | 132 |
| Bat | ND | 2001 | 2110 | Hic | ND | 1122 | 929 |
|  | HF | 2489 | 2162 |  | HF | 1177 | 1099 |
| | ND ∩ HF | 259 | 336 | | ND ∩ HF | 102 | 99 |
| Muscle | ND | 933 | 1448 | 7 tissues | ND | 3 | 4 |
|  | HF | 1491 | 917 |  | HF | 5 | 3 |
| | ND ∩ HF | 203 | 122 | | ND ∩ HF | 3 | 3 |

DEGs were screened using the criteria of $p < 0.05$ and fold change (FC) $> 1$ . $J > N$ , the expression level in B6J was higher than in B6N; $J < N$ , the expression level in B6J was lower than in B6N; $ND \cap HF$ , the number of overlapping DEGs among ND-fed and HF-fed mice. iWAT, inguinal white adipose tissue; eWAT, epididymal white adipose tissue; BAT, brown adipose tissue; Muscle, skeletal muscle; Hyt, hypothalamus; Hic, hippocampus. ND, normal diet; HF, high-fat diet.

**Fig 6.**
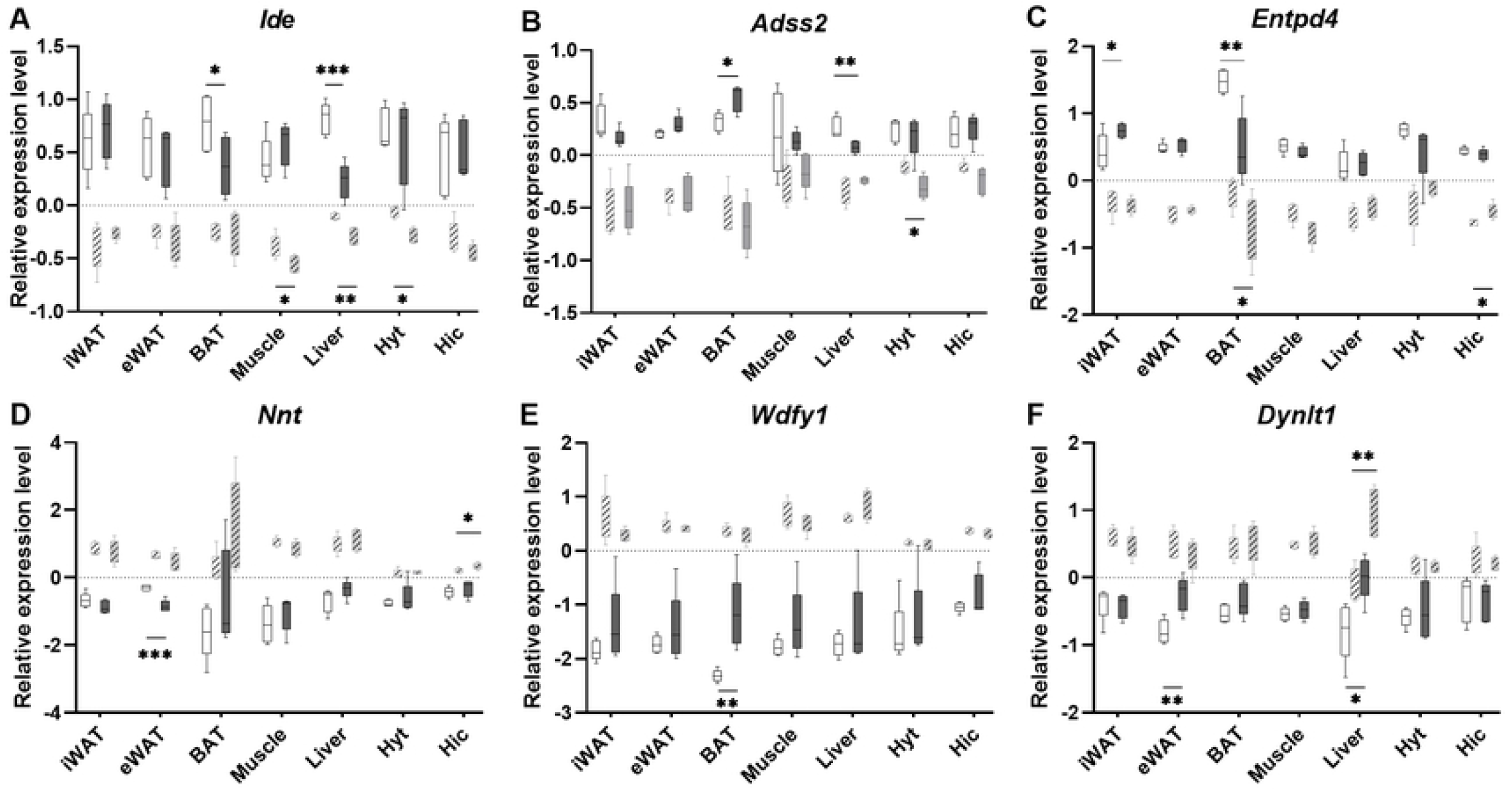
Expression levels of DEGs between B6J and B6N on ND or HF. The gene name is indicated at the top of each plot, and the y-axis represents the normalized signal values. (A) *Ide*, insulin degrading enzyme; (B) *Adss2*, adenylosuccinate synthase 2; (C) *Entpd4*, ectonucleotide triphosphate diphosphohydrolase 4; (D) *Nnt*, nicotinamide nucleotide transhydrogenase; (E) *Wdfy1*, WD repeat and FYVE domain containing 1; (F) *Dynlt1*, dynein light chain Tctex-type1. ND, normal diet; HF, high-fat diet. Open boxes, ND-fed B6J; hatched boxes, ND-fed B6N; filled boxes, HF-fed B6J; filled and hatched boxes, HF-fed B6N. Asterisks (*) indicate significant differences (\**p*<0.05, \*\**p*<0.01) between ND and HF.

The effect of a high-fat diet on the expression of these overlapping genes was not observed in most tissues. However, decreased *Ide* expression and increased *Dynlt1* expression in the liver (Fig 6A and 6F), and decreased *Entpd4* expression in BAT (Fig 6C) were observed under HF conditions compared to those observed under ND conditions.

## Discussion

Phenotypic differences among B6 substrains have been reported in many studies in a variety of fields, including metabolism [7, 12, 14-19, 31, 33], alcohol preference [35-38], stress response [27, 32, 39-44], cardiovascular [20, 21, 45], oxidative stress [22,23], vision [8-10], bone [28], liver [29, 30, 46], kidney [24, 47, 48], microbiota [15], and seizures [49]. However, the genes responsible for these phenotypic differences have not been identified. Often, NNT deficiency due to a spontaneous missense mutation of *Nnt* in B6J mice is considered to be responsible for these phenotypic differences and abnormal traits in B6J mice [11-24], including metabolic dysfunctions such as impaired glucose tolerance, diminished insulin secretion, and overweightness. This speculation is based on the fact that the enzymatic function of NNT is to pump protons across the inner mitochondrial membrane [26]. However, the difference in the phenotype between B6J and B6N is not always correlated to the absence or presence of NNT. For example, *Nnt*-deficient B6J mice (S1 Fig) in the present study were significantly underweight compared to B6N mice (Fig 1), as in previous studies [15, 29-32]. However, there are conflicting data on body weight differences: overweight B6J mice [12-14, 27, 28] or unchanged weight [16, 17, 33]. Similarly, there are also conflicting data on the differences in blood insulin and glucose levels between B6J and B6N mice with several studies reporting that B6J mice have lower insulin levels [12, 14, 17-19, 29, 31], higher glucose levels [12, 15, 18, 19, 31], and lower glucose levels [14] compared to B6N mice, whereas some studies did not report any differences in glucose [16, 17, 29, 31, 33] or insulin [16, 31]. In our study, we observed higher glucose levels but lower insulin levels in *Nnt*-deficient B6J mice compared to *Nnt*-wild type B6N mice (Fig 4). Given the discrepancy between the B6J and B6N phenotypes and NNT levels, we speculate that genetic mutations other than *Nnt* mutation alleles are present in either B6J or B6N.

We investigated DEGs in iWAT, eWAT, BAT, muscle, liver, hypothalamus, and hippocampus tissues between B6J and B6N, and identified six DEGs, including *Nnt*, which overlapped in all seven tissues regardless of dietary conditions (Fig 6). Three DEGs, *Ide, Adss2*, and *Entpd4*, were significantly more expressed in B6J mice than in B6N mice, whereas three DEGs, *Nnt, Wdfy1*, and *Dynlt1*, were significantly less expressed in B6J mice than in B6N mice.

Previously, *Wdfy1* and *Entpd4*, have been reported to be differentially expressed in the brain and pancreas between B6J and B6N [37, 50],; thus, our findings are consistent with those of this study. High *Wdfy1* expression in the brain of B6N is speculated to be related to reduced alcohol intake in B6N mice [37], whereas low *Wdfy1* expression in the pancreas of B6J mice is associated with the progression of chronic pancreatitis in B6J mice [50]. Although these previous studies did not focus on *Entpd4* [37, 50], its higher expression in B6J mice compared to B6N mice is consistent with the finding of the present study.

Watkins-Chow and Pavan have reported that the presence of increased copy number variation (CNV) at the *Ide* locus in B6J mice results in increased *Ide* expression [51]. The CNV may contribute to the divergence in *Ide* expression between B6J and B6N observed in this study. The lists of B6N substrains of protein-inactivating sequence variations (sequence variations causing premature stop codons, loss of stop codons and SNPs, and short in-frame insertions and deletions) that referenced the B6J mouse genome [52] did not include the *Ide, Adss2, Entpd4, Wdfy1*, and *Dynlt1* genes identified in this study. Therefore, CNVs in B6J and B6N mice should be investigated in addition to indels and SNPs [53]. Additionally, IDE is an enzyme that degrades insulin. Considering the lower blood insulin concentration and slightly higher glucose concentration in B6J mice compared to B6N mice (Fig 4A and 4H), the higher *Ide* expression level in B6J mice is consistent with the lower blood insulin concentration (Fig 6A). These results suggest that *Ide* may be responsible for the phenotypic differences in insulin secretion between B6J and B6N mice. *Ide* also plays a role in type 2 diabetes and Alzheimer’s disease [54]; thus, caution should be exercised when using B6 substrains in studies targeting these diseases. Nevertheless, genetic differences among substrains can be a powerful tool for studying loci that influence disease-related phenotypes. Therefore, one approach to Alzheimer’s disease research would be to compare traits associated with Alzheimer’s in the B6J substrain, which has high *Ide* expression, and the B6N substrain, which has low *Ide* expression.

A previous study has reported that mice lacking nucleobindin 2 (*NUCB2*), a precursor of nesfatin involved in appetite regulation, exhibit insulin resistance and high *Wdfy1* expression in their visceral macrophages [55], suggesting that *Wdfy1* may be also involved in insulin signaling. However, since the genetic background of the floxed mice used in the *NUCB2* paper [55] was B6J, and that of the crossbred recombinase mice was “C57BL/6J; C57BL/6N” [56], we cannot rule out the possibility that high *Wdfy1* expression was simply due to comparing B6N, which has high *Wdfy1* expression, with B6J, which has low expression. Similarly, the use of mice with a mixed background of B6J and B6N in immunological studies leads to confounding result interpretations. For example, the effect of granzyme A (GZMA) on viral arthritis remains inconclusive because the phenotypes observed in experiments using *GZMA* knockout mice vary from paper to paper [57]. However, the genetic background of *GZMA* knockout mice was found to be a mix of B6J and B6N, indicating that viral arthritis ameliorated in *GZMA* knockout mice was not the consequence of loss of GZMA expression, but rather of the genetic background of B6N, including *Nnt* [57]. The importance of substrain selection has also been reported in studies on COVID-19, where the B6J substrain was reported to be unsuitable for studies on severe COVID-19 or COVID-19-related long-term lung diseases because of its low infectivity [58].

To the best of our knowledge, no literature on *Adss2* and *Dynlt1* as DEGs between B6J and B6N mice currently exists. Nevertheless, we have identified DEGs between B6J and B6N (S1 Table), which might help plan mouse experiments. Our findings may be useful for revisiting past studies that have used B6J, B6N, or B6 substrains to determine whether their results answer the original research objectives or are merely a measurement of the differences between B6J and B6N.

## Conclusion

*Nnt* mutations in B6J mice have been implicated as a cause of obesity and various metabolic abnormalities and described as the causative genetic explanation for the phenotypic differences between B6J and B6N mice. However, there is a discrepancy in metabolic traits between B6J and B6N that cannot be explained by *Nnt* alone, raising the possibility that mutant alleles other than *Nnt* exist. We identified genes such as *Ide, Adss2, Entpd4, Wdfy1*, and *Dynlt1* that are differentially expressed between B6J and B6N mice. Since both B6J and B6N have been widely used to generate genetically modified mice as background strains in medical biology experiments, it is important to be aware of the genetic differences between the two to avoid misinterpretation of the experimental results.

## Supporting Information

**S1 Table. Differentially expressed genes between inguinal white adipose tissue, epididymal white adipose tissue, brown adipose tissue, skeletal muscle, liver, hypothalamus, and hippocampus of B6J and B6N mice fed a normal diet or high-fat diet**. The *p-*value was calculated using the moderated *t*-test, and the false discovery rate was calculated using the Benjamini-Hochberg method. DEGs, differentially expressed genes; FC, fold change: FDR, false discovery rate.

**S1 Fig. Genotyping of the *Nnt* gene**. Polymerase chain reaction analysis of *Nnt* alleles. DNA was obtained from the tail of B6J and B6N mice. The amplification products were 579 bp and 743 bp for the wild-type and mutant alleles, respectively.

